# Onset of sexual maturity of sexually propagated and wild *Favites abdita* colonies in northwestern Philippines

**DOI:** 10.1101/2021.01.06.425549

**Authors:** KG Bonilla, JR Guest, DW dela Cruz, MV Baria-Rodriguez

## Abstract

Scleractinian corals are modular colonial organisms and are the main framework builders of coral reefs. Most corals reproduce by broadcast spawning with external fertilization and these processes are essential to replenish reef coral populations. Despite decades of research, many aspects of coral reproductive biology remain poorly studied. For example, two important reproductive life history traits, colony size and age at the onset of sexual maturity, are mostly unknown for many reef-building corals. In this study, wild colonies of different size classes and colonies of a known age (i.e. colonies sexually cultured and reared from larvae to adults) of the massive scleractinian *Favites abdita* were examined for the presence or absence of mature oocytes to determine size and age at the onset of sexual reproduction. Fecundity at the onset of reproductive maturity across size classes of wild colonies was also determined. Surveyed and sampled colonies were grouped into three size classes based on maximum colony diameter (A = 0.1-4.0 cm, B = 4.1-8.0 cm, and C = >8.1 cm). For both wild and sexually propagated colonies, the smallest colonies containing gametes were 1.8 cm in diameter, suggesting that this is the minimum colony size at onset of sexual maturity. Colonies of size class A had lower mean oocyte counts per polyp (44 ± 6.08; mean ± SE) compared to colonies of classes B and C (469 ± 62.41, 278 ± 57.15, respectively). However, mean oocyte geometric mean diameter of size class A colonies was greater (340.38 ± 7.68 μm) than classes B and C (283.96 ± 6.94 μm, 317.57 ± 9.18 μm, respectively). Results of this study bring in to question the widely applied operational definition of coral juveniles being colonies ≤4.0 cm diameter and suggest that even quite small colonies may play a role in contributing to the natural larval pool on reefs than previously thought.

## 1. Introduction

Scleractinian corals are the main framework builders on coral reefs, creating much of the structural complexity needed to support species biodiversity (Moberg and Folke 1999). Corals are modular organisms that can either reproduce asexually through fragmentation or sexually through the production of gametes and larvae (Harrison and Wallace 1990). Mature gametes are either brooded internally wherein fertilization takes place inside the polyp and competent larvae are released into the water column, or gametes are broadcast spawned into the water column for external fertilization (Harrison and Wallace 1990, Baird et al. 2009, Schmidt-Roach et al. 2012). Hard corals have a biphasic life cycle having a short larval phase, followed by a longer benthic phase until sexual maturity (Harrison 2011). Larvae settle and metamorphose on hard substratum within a few days after fertilization, then calcify and grow to adult until it reach sexual maturity (Miller and Mundy 2003; Nishikawa et al. 2003; Toh et al. 2012).

Coral juveniles are often operationally defined as sexually immature coral colonies with a diameter of less than or equal to 4 cm (Bak and Engel 1979; Harrison and Wallace 1990, Soong 1991; Edmunds 2007). This definition has been used to describe juveniles of different coral species with various life-history strategies in numerous studies (Kojis and Quinn 1982; Szmant 1986; Hall and Hughes 1996; Bak and Meester 1999; Edmunds and Carpenter 2001; Kai and Sakai 2008). During reef surveys, classification and differentiation of juveniles and reproductively mature adults is usually based on colony size only. However, reproductive maturity of adult corals is a function of polyp age, colony age, and colony size (Connell 1973; Kojis and Quinn 1985; Harrison and Wallace 1990; Kai and Sakai 2008). For example, the massive species *Coelastrea aspera* was described to have the capability to allocate resources to reproduction upon reaching a certain colony age regardless of colony size (Kai and Sakai 2008). In the case of *Favites chinensis* and *Goniastrea favulus*, colony size, determined the start of sexual maturity while colony age influenced polyp fecundity (with older colonies having greater fecundity than younger colonies) (Kojis and Quinn 1985, Kai and Sakai 2008). Hence, once a certain colony size or age has been achieved, polyps will begin to produce gametes (Kojis and Quinn 1985; Szmant-Froelich 1985, Kai and Sakai 2008).

Age/size at onset of sexual maturity of corals and fecundity (average egg count per polyp) are traits that have important implications for coral population dynamics and demography (Lukoschek et al. 2013; Adjeroud et al. 2018). In addition, coral fecundity is a good indicator of the coral’s health, energy invested in reproduction, and energy allocated to offspring (Bell 1980; Eckelbarger 1986; McGinley et al. 1987). Available information on the size at maturity of branching corals come mostly from fast-growing *Acropora* species found in Australia, Japan, and the Philippines (Van Moorsel 1988, Wallace 1985, Kojis 1986, Iwao et al. 2010, Baria et al. 2012, dela Cruz and Harrison 2017). While sizes at maturity of submassive and massive corals are known for the species from the genera, *Favia, Coelastrea*, and *Goniastrea*, in Australia, Panama, Puerto Rico, and Japan (Babcock 1984, Szmant-Froelich 1985, Kojis and Quinn 1985, Babcock 1986, Sakai 1998b, Soong 1993). Slow-growing massive species are important reef builders and are more stress-tolerant (Darling et al. 2012). Despite the importance of massive species on the reef, there are relatively few studies on their colony size and age at the onset of sexual maturity. Moreover, previous studies that reported the colony sizes at maturity of some massive species were not able to establish the age at reproductive maturity due to varying growth rates.

The purpose of this study is to determine the age and colony size at the onset of sexual maturity of *Favites abdita* and compare fecundity among size classes. Information on the colony size and age at first reproduction of most slow-growing massive species as well as the fecundity across different size classes are mostly lacking despite their important implications on coral population dynamics and demography. In the present study, the maximum colony diameters of sexually produced and wild *F. abdita* colonies in northwestern Philippines were measured and were examined for the presence of mature oocytes. Both wild *F. abdita* colonies and colonies of a known age were surveyed for the first time in this study to determine the colony size and age at the onset of sexual maturity (i.e. smallest colony containing mature oocytes). Samples were also collected from wild *F. abdita* colonies to determine the fecundity across different size classes and to help infer how much do small colonies (≤4 cm maximum diameter) contribute to the larval pool on the reef.

## 2. Materials and methods

The massive *F. abdita* is a hermaphroditic broadcast spawner, commonly found at a depth range of 1 to 15 m in Bolinao-Anda Reef Complex and is known to spawn every May (Maboloc et al. 2016). In this study, the onset of reproductive maturity was characterized by the presence of mature oocytes *in situ* (Wallace 1985; Sakai 1998a, 1998b; Llodra 2002; dela Cruz and Harrison 2017). Colony diameters of sexually propagated *F. abdita* and wild colonies were measured and colonies were examined for the presence and absence of mature oocytes. In addition, wild colonies sampled from the reef were dissected to determine the fecundity (oocyte count, oocyte size, and oocyte volume per polyp) across different size classes. Surveyed sexually propagated *F. abdita* and sampled wild colonies were grouped into three size classes based on maximum diameter as follows: A = 0.1-4.0 cm, B = 4.1-8.0 cm, and C = >8.1 cm. Size class A is the size range of corals that are operationally classified as juveniles (less than or equal to 4.0 cm maximum diameter) in most studies, while size classes B and C are the sizes typically considered to be adults (Kojis and Quinn 1982; Szmant 1986; Hall and Hughes 1996; Bak and Meester 1999; Edmunds and Carpenter 2001; Kai and Sakai 2008).

### 2.1 Sexual propagation, nursery rearing, and transplantation of *F. abdita* colonies

Colonies of known ages, that were sampled in the present study, were grown from fertilized eggs spawned in 2009 as part of a different project to examine the technical feasibility of coral reef rehabilitation using sexual propagation (Guest et al. 2010; 2014). Full details of the culturing techniques, the method for rearing juvenile corals in nurseries, and the protocol used for transplantation are described in Guest et al. (2010; 2014) and Toh (2014) and are briefly summarized below. Seven gravid *F. abdita* colonies with a geometric mean diameters of 19.1 ± 6.4 cm (mean ± SD) were collected from the reefs close to the Bolinao Marine Laboratory (BML), University of the Philippines (16°44’N, 119°94’E) on 13 May 2009, 4 days after the full moon. Colonies were placed in 400 L rectangular plastic tanks with flow-through seawater at the hatchery facility of BML and were monitored for spawning activity. All spawning occurred on May 14 (5 nights after full moon on 9 May) between the hours of 20:00 and 21:00 h. Gamete bundles were collected, transferred to 60 L plastic tanks with UV-treated 1 μm filtered seawater (FSW), and were gently agitated to separate gametes for cross-fertilization. Embryos were washed of excess sperm and were transferred to three 400-900 L rearing tanks with FSW.

Reared larvae were settled onto biologically conditioned “plug-ins” consisting of a polyethylene wall plug embedded in a cylindrical cement head (20 mm diameter, 15 mm height, 1,492 mm^2^ surface area) (Guest et al 2014). At three days post-fertilization, plug-ins were placed at the bottom of the rearing tanks and were left for 10 days to permit coral larval settlement and metamorphosis with daily change of 50-100% FSW. Plug-ins with at least one settled coral were inserted into 1-m^2^ rearing trays made of polyethylene mesh with 1 cm diameter holes and PVC pipe frames. Trays were placed in 400 L cement tanks with flowthrough seawater for 12 months of *ex situ* rearing. After 12 months, in May 2010, all plug-ins with a living coral were transferred to an *in situ* nursery in Malilnep channel (16°26’N, 119°56’E) consisting of two fixed tables (2.5 m long, 0.6 m wide, 0.8 m high) constructed using angle-iron bars hammered into the substrate at 1.8 m depth. After a further 13 months at the *in situ* nursery (25 months post-fertilization), plug-ins (n=120) were outplanted in Lucero, Marcos, and Caniogan reefs (Figure 1) in June 2011.

**Figure 1.**
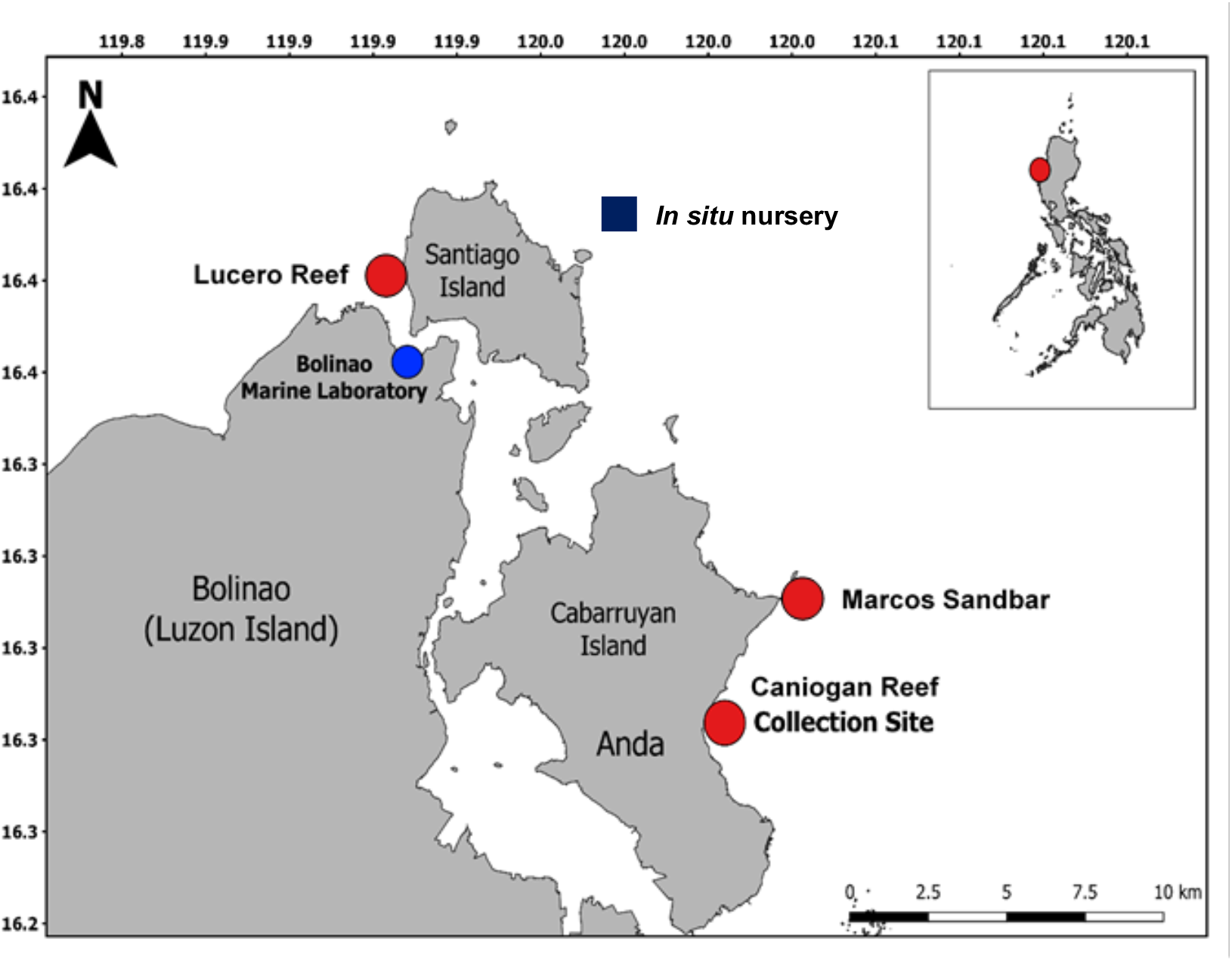
Site map showing the *in situ* nursery, the transplant sites (Lucero Reef, Marcos Sandbar, Caniogan Reef) of sexually cultured and reared *F. abdita* colonies, and the collection site (Caniogan Reef) of wild *F. abdita* samples.

### 2.2 Field survey of sexually propagated *F. abdita* transplants

To determine the size at the onset of reproductive maturity (i.e. smallest colony with mature oocytes) of sexually propagated *F. abdita* transplants, as well as to compute for the proportion of gravid colonies per size class, all surviving transplanted colonies were surveyed and sampled for presence or absence of visible mature oocytes 7 to 9 days before the full moon of May 2015 (71 months post-fertilization). The maximum diameter of colonies was measured using a caliper and maturity was assessed by carefully breaking one or several polyps using hammer and chisel. The presence or absence of mature oocytes (pink to light red in color) was noted in each of the colony.

### 2.3 Survey and sampling of wild *F. abdita* colonies

#### 2.3.1 Collection of samples

Collection of samples was carried out on 19 to 20 April 2018, 39 to 40 days before the full moon of May 2018. A total of 31 wild *F. abdita* colonies found at depths between 2 and 6 m in Caniogan (16°18’N, 120°01’E; Figure 1) were measured with a caliper and were sampled using a hammer and a chisel to determine the size at the onset of sexual maturity and fecundity. Sampled colonies were >1 m apart to minimize the possibility of collecting samples from clonal colonies. Fragments (<1–5 cm) were collected from the center of the colonies to avoid sampling the sterile polyps located at colony edges (Hall and Hughes 1996, Sakai 2008b). For colonies <5 cm diameter, the entire colony was carefully removed and collected. Samples were then placed in pre-labeled resealable containers with seawater and were transported to the laboratory for tissue processing.

#### 2.3.2 Fixation and decalcification of coral samples

Samples were fixed with Zenker’s Solution (50 g zinc chloride and 25 g potassium dichromate per liter of filtered seawater with 50 ml of 37% formalin per liter of seawater) for 24 hours, then rinsed with running filtered seawater for 24-hours before being decalcified in 10% buffered HCl solution (0.7 g EDTA, 0.008 g sodium potassium tartrate, 0.14 g sodium tartrate, and 100 ml HCl in 900 ml distilled water). The acid solution was replaced every 2 to 3 days until samples are completely decalcified. Samples were rinsed with running fresh water for an hour and were stored in 70% ethanol until dissection and histological processing. (Szmant-Froelich et al. 1985 and Glynn et al. 1991).

#### 2.3.3 Dissection and histological slide examination

Fixed coral tissue samples were dissected on a petri dish using fine needles under a dissecting microscope. Ten polyps were randomly chosen from each sample to check for the presence or absence of oocytes and to determine the corresponding fecundity of samples with oocytes. The total oocytes per polyp were counted and ten randomly selected oocytes were measured. Maximum diameter (*D1*) and maximum perpendicular diameter (*D2*) were measured using the Motic software version 3.0 to calculate the geometric mean diameter (GMD) using the formula 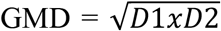. Oocyte volume (V) was also taken using the spherical formula 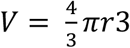 where 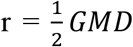. The total volume of oocyte per polyp was calculated by multiplying the mean oocyte number per polyp by the mean oocyte volume (Hall and Hughes 1996). Representative samples (n=14) were dehydrated, cleared, and embedded in paraffin for histological analysis. Latitudinal cross-sections of polyps at 3-4 μm thick were mounted onto glass slides and were stained with hematoxylin and eosin. Prepared slides were examined under a compound light microscope and oocytes were categorized either as Stage I, II, III, IV, or V based on Szmant-Froelich et al. (1985) and Shikina et al. (2013).

### 2.4 Statistical analysis

Data did not meet the parametric test assumptions of Analysis of Variance (ANOVA). Thus, differences in oocyte count, oocyte GMD, and total oocyte volume per polyp across size classes were analyzed using the nonparametric Kruskal-Wallis test and Dunn’s multiple comparison test for post-hoc. Samples that were non-gravid were not included in the report of fecundity across different size classes to avoid underestimation.

## 3. Results

A total of 98 sexually produced and transplanted *F. abdita* colonies were examined to check for the presence or absence of oocytes, while 31 wild colonies were sampled to determine fecundity across size classes and minimum colony size at the onset of sexual maturity. Among the 98 transplanted 6-yr old colonies, 10 colonies were assigned to size class A, 66 colonies were in size class B and 22 colonies were in size class C. Of the 31 wild colonies, 18 colonies were in size class A, while 9 and 4 colonies respectively were in size classes B and C.

The smallest diameter of gravid colonies measured for both sexually produced and wild colonies was 1.8 cm (Figure 2A). The proportion of gravid sexually produced and wild *F. abdita* colonies increased with colony size (Figure 3), with >40% of both transplanted and wild colonies mature at size class B, and >70% of colonies mature at size class C (Fig. 3). Fecundity of wild colonies varied between size classes. Size class A colonies had an average fecundity of 44 ± 6.08 (mean ± SE) oocytes per polyp, whereas size classes B and C colonies had average fecundities of 469 ±62.41 and 278 ± 57.15 oocytes per polyp respectively (Figure 4A). Mean oocyte GMD across size classes ranged from 283.96 ± 7.68 to 340.38 ± 9.18 μm (Figure 4B). The total volume of oocytes was smaller in size class A with 0.93 ± 0.34 mm^3^ per polyp compared to size class B and C with 3.78 ± 3.10 and 7.06 ± 2.16 mm^3^ per polyp respectively (Figure 4C). Number (H = 22.842, 2 d.f., p = 0.00001) and sizes of oocytes (H = 20.254, 2 d.f., p = 0.00004) per polyp were significantly different across size classes. Total oocyte volume per polyp, however, was not significantly different across size classes (H = 2.1115, 2 d.f., p = 0.3479). Moreover, most of the oocytes of wild colonies were at stages III to IV while observed spermaries were at stage IV (Figure 5 and Table 1).

**Figure 2.**
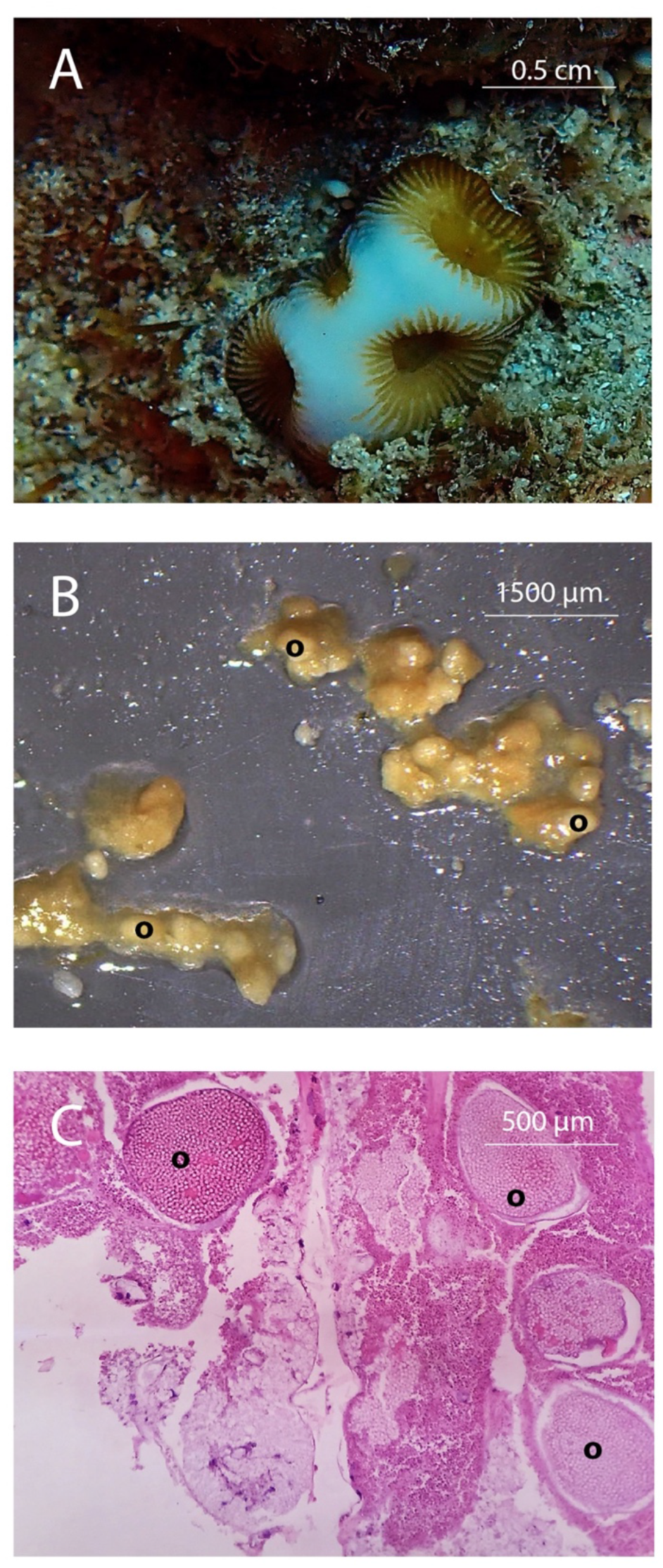
Wild *F. abdita* colony (1.8 cm diameter) (A) with visible oocytes seen during dissection (B) and histological slide examination (C). O = oocytes

**Figure 3.**
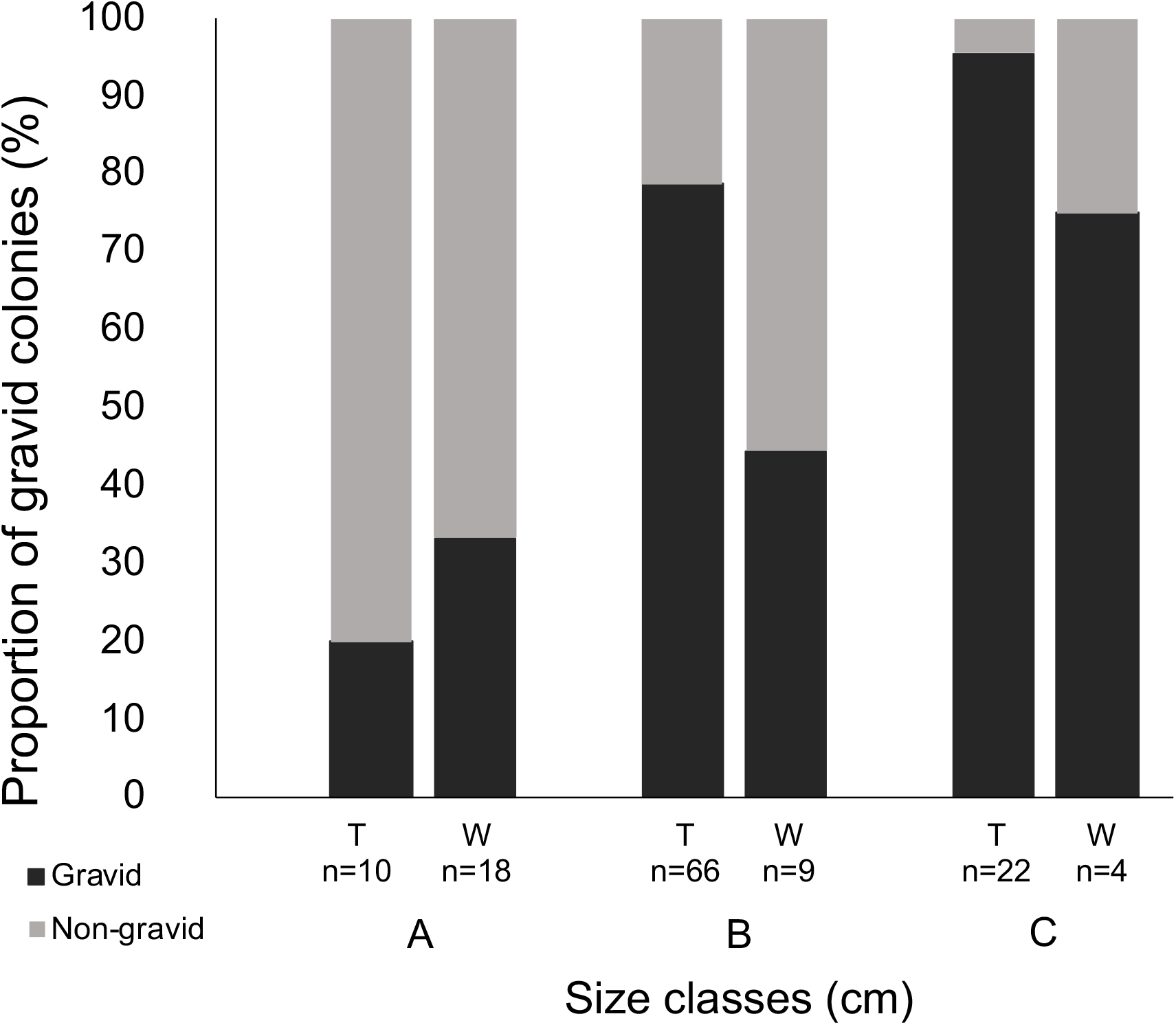
Proportion (%) of gravid and non-gravid colonies of *F. abdita* across size classes (A = 0.1-4.0 cm, B = 4.1-8.0 cm, and C = >8.1 cm maximum diameter). Transplant (T) data represent the presence and absence of mature oocytes of sexually cultured corals in the hatchery and outplanted on the reef while wild (W) data represent the presence and absence of mature oocytes of natural colonies found on the reef, n = total number of colonies examined.

**Figure 4.**
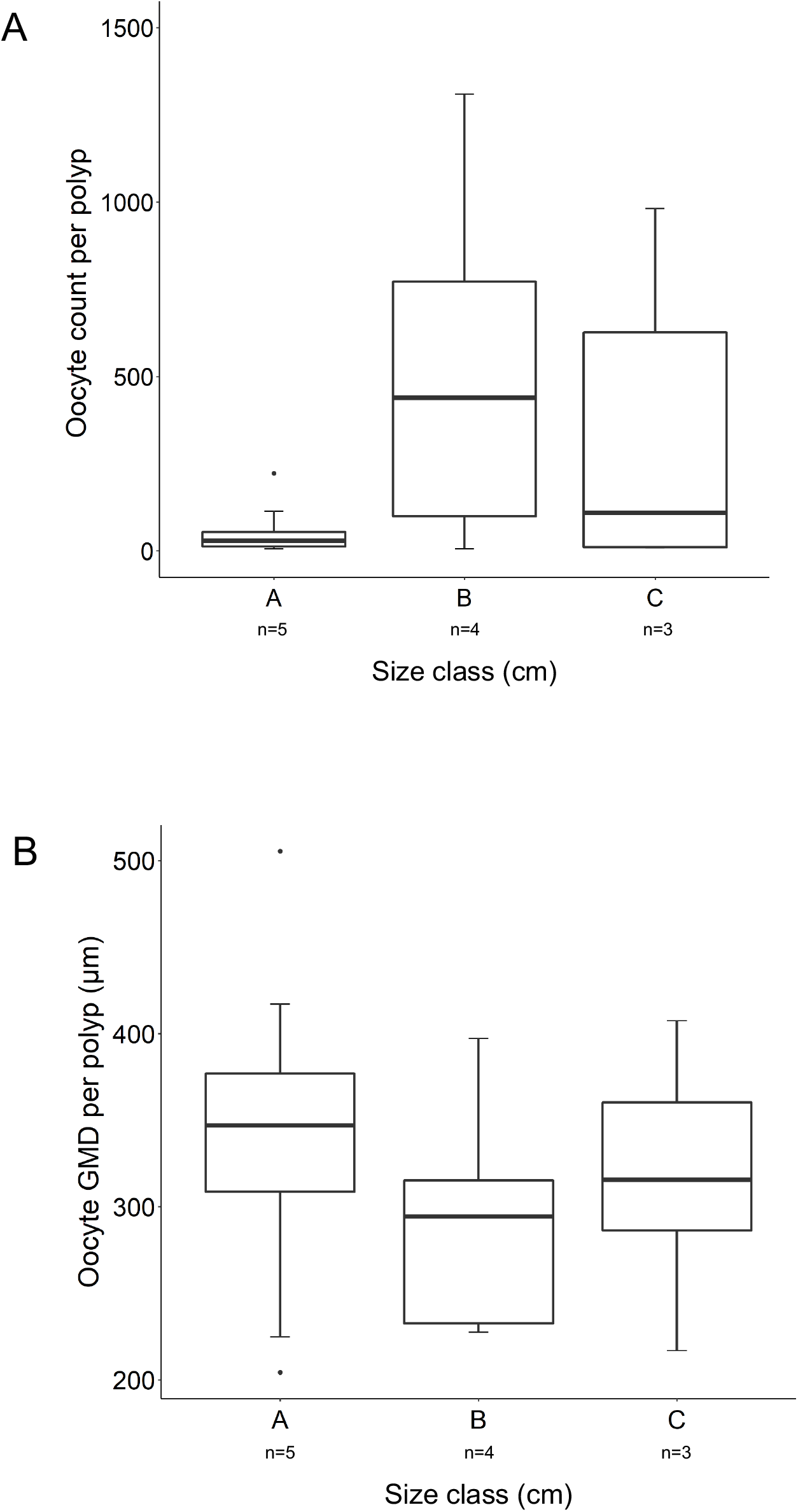
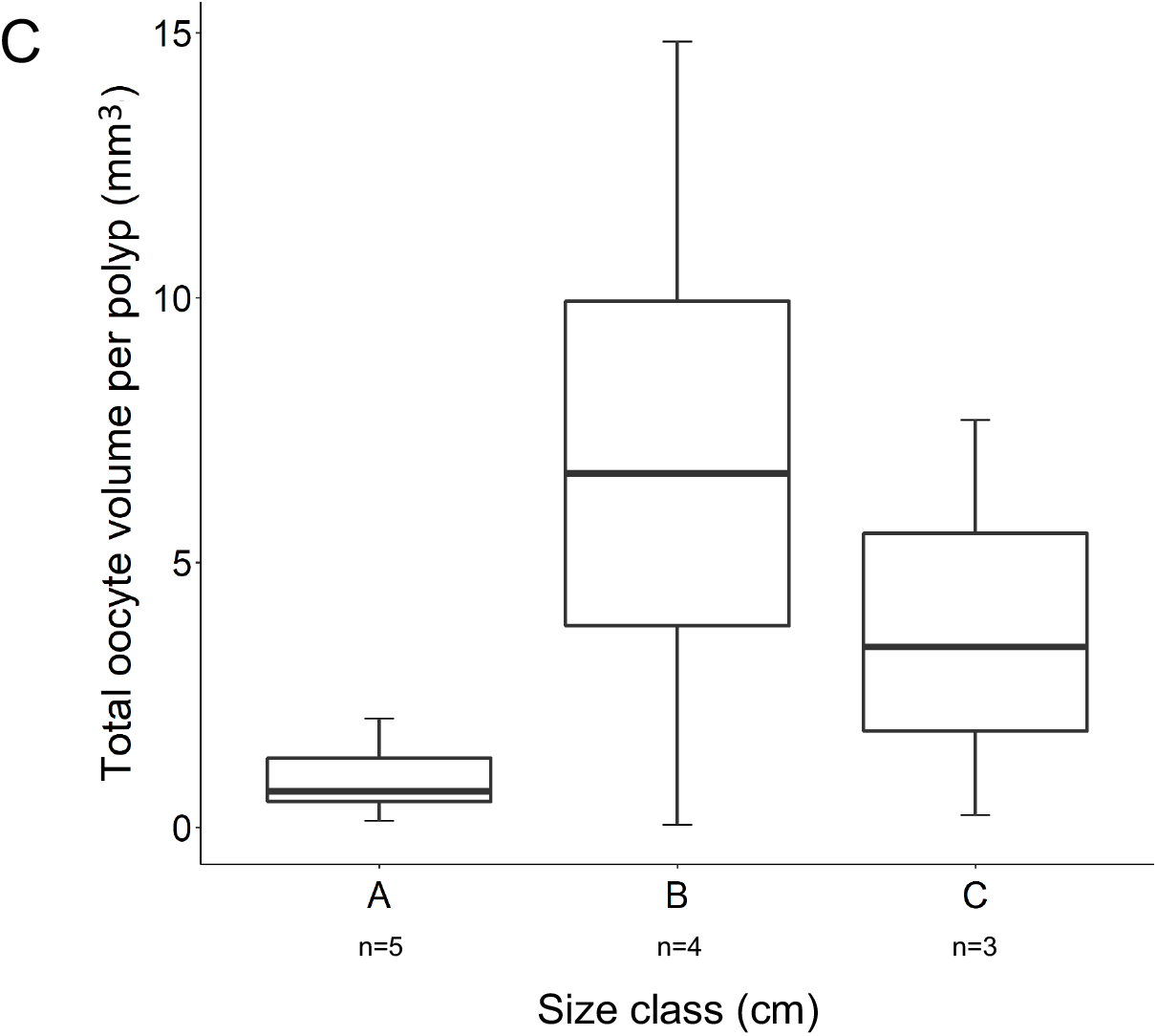
Oocyte count (A), oocyte geometric mean diameter (μm) (B), and total oocyte volume (mm^3^) (C) per polyp across size classes (A = 0.1-4.0 cm, B = 4.1-8.0 cm, and C = >8.1 cm maximum diameter) of gravid wild *F. abdita* colonies. n = number of colonies sampled.

**Figure 5.**
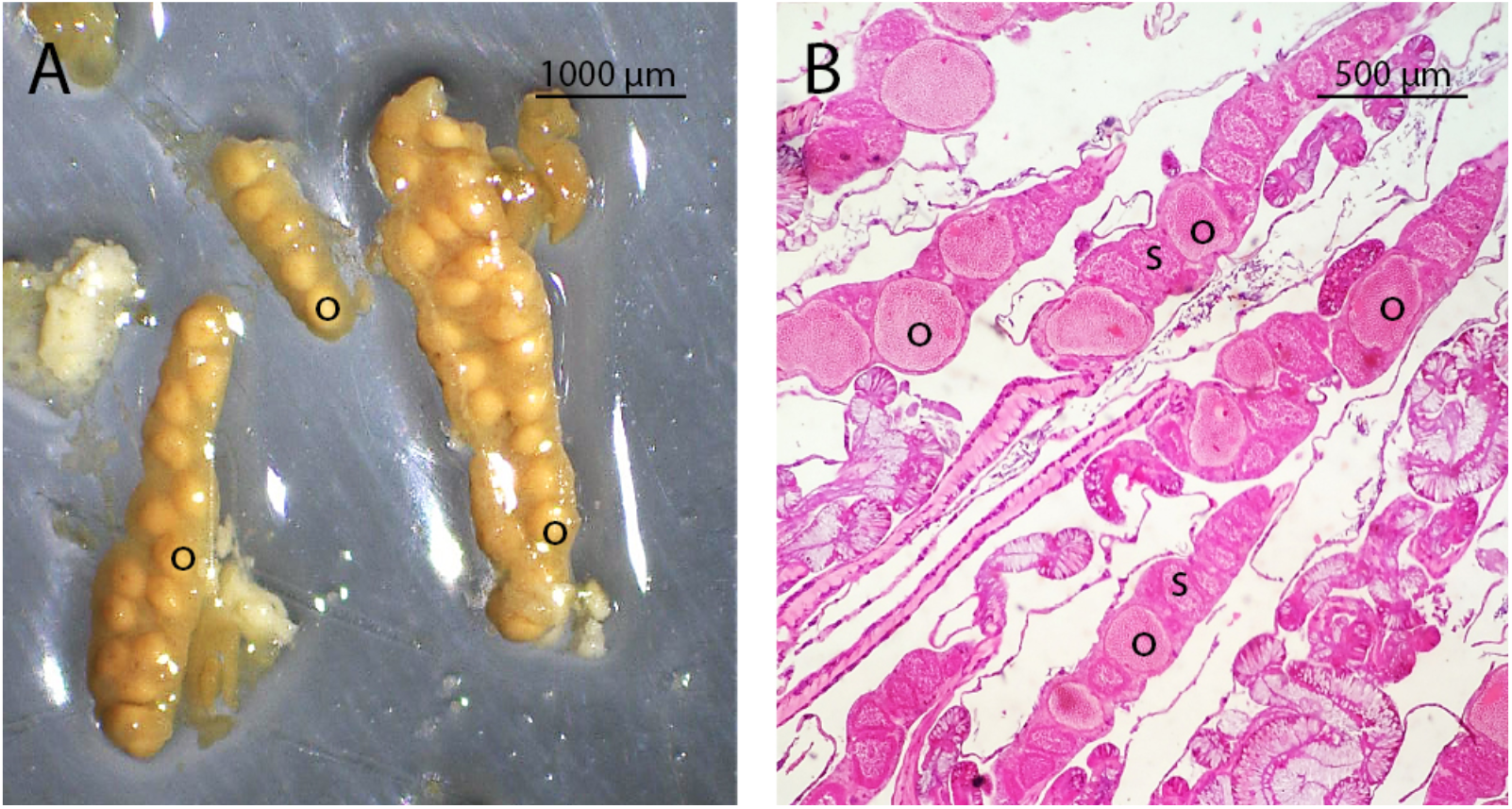
Strings of gametes under a dissecting microscope (A) and a histological slide showing stage IV oocytes and spermaries (B). O = oocytes, s = spermaries.

**Table 1.**
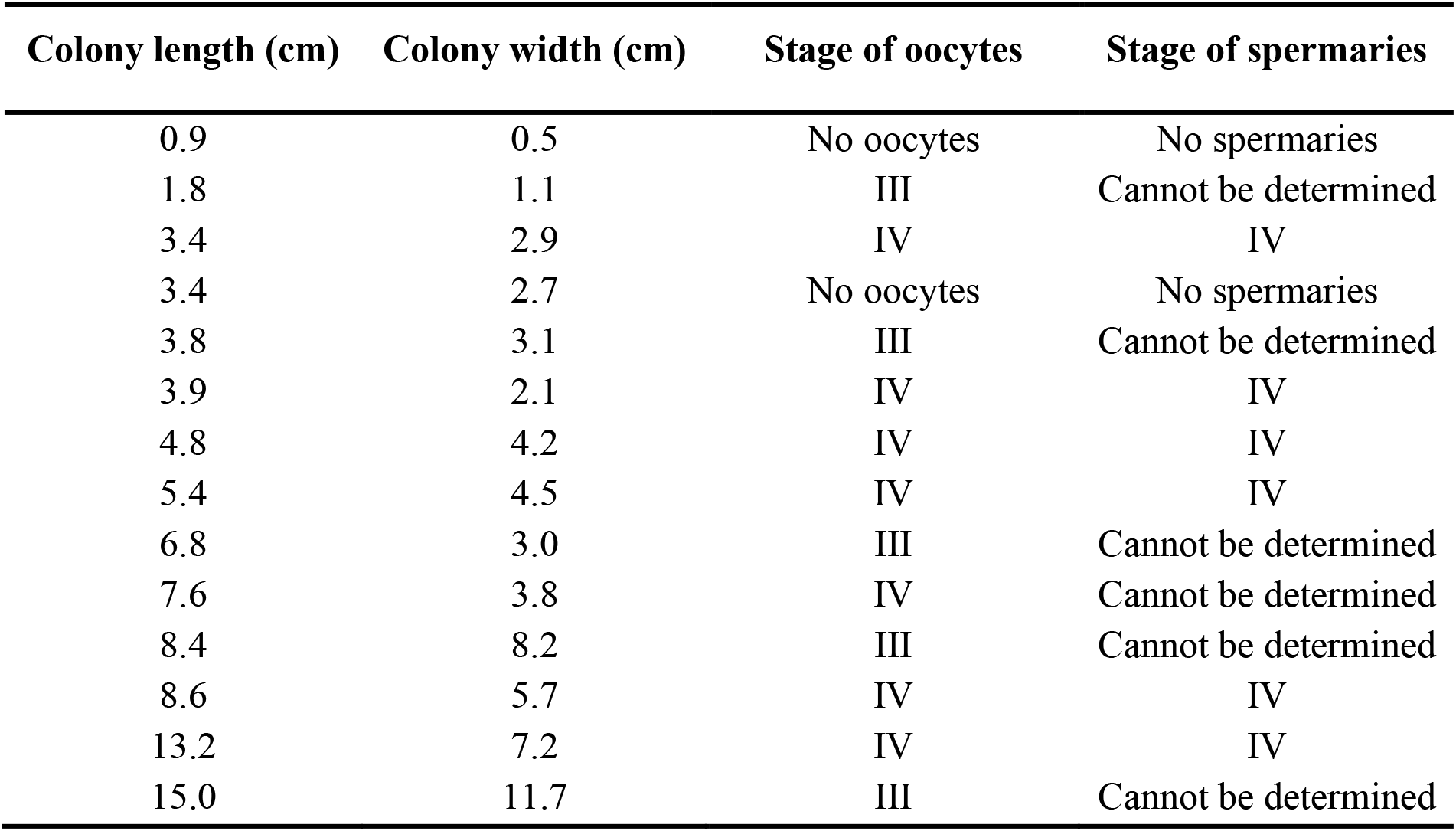
Representative colonies sampled for histological preparation and the corresponding gamete stages upon examination.

## 4. Discussion

The age and colony size at onset of sexual maturity of common reef-building corals have received less research attention to date compared to other aspects of coral reproductive biology and ecology such as reproductive timing, spawning, and larval culture (Villanueva et al. 2008; Baria et al. 2012; Guest et al. 2014). This is the first study to determine the size at the onset of sexual reproduction of slow-growing *F. abdita* in both transplanted colonies of a known age and wild colonies of different size classes. Both wild colonies and sexually produced colonies of known age were reproductive at relatively small sizes (i.e., < 2 cm diameter). This research suggests that small colonies, that are usually defined as juveniles during reef surveys (< 4 cm), may be reproductively mature colonies, at least for massive and slow-growing corals. The implication of these findings is that relatively small coral colonies may contribute to replenishment of coral populations via sexual reproduction.

For both sexually produced (~6 year-olds) and wild colonies, the smallest colony containing oocytes was 1.8 cm diameter. Corals are modular organisms that can undergo fission and fusion. It is possible, therefore, that smaller natural colonies that resembled juveniles, may actually have resulted from remnants of larger, older colonies. This was not the case for the smallest reproductively mature colony of sexually propagated *F. abdita* transplants, as these colonies had been tagged and their exact ages were known. Our results support previous studies of other massive species that found small colonies (i.e., <4 cm diameter) to contain gametes. For example, the smallest mature colonies of *Favia fragum, Coelastrea aspera* and *G. favulus* range in size from 1.6 to 3 cm (Babcock 1984, Kojis and Quinn 1985, Wallace 1985, Soong 1993, Sakai 1998b).

The number of oocytes per polyp in wild *F. abidta* colonies increased as colony size increased from size class A (0.1-4.0 cm) to B (4.1-8.0 cm) with mean oocyte counts per polyp increasing from 44 ± 6.08 to 469 ± 62.41 (mean ± SE). An increase in fecundity with increasing colony size has also been observed for other slow-growing corals, such as *G. aspera* (Babcock 1984). Colonies of *G. aspera* with a diameter range of approximately 0.1 to 4.0 cm had no oocytes while 4.2 to 8.0 cm colonies had 47,500 oocytes per colony which increased to 215,400 oocytes as colonies reached diameters of 8.2 to 12.0 cm (Babcock 1984). The average number of oocytes per polyp, however, decreased to 278 ± 57.15 (mean ± SE) for larger colonies in size class C, >8.1 cm diameter. Smaller colonies were also observed have a lower relative volume of female gametes per polyp than larger colonies. This strategy channels the energy more to growth and survival of small colonies as eggs are lipid-rich and are energetically costly to produce (Kojis 1982; Szmant 1986; Hall and Hughes 1996; Sakai 1998a; Sakai 1998b; Kai and Sakai 2008). Moreover, producing large quantities of small oocytes in reefs, where reproducing colonies are sparsely distributed may be advantageous as this strategy increases chances of successful fertilization (Babcock 1984).

Histological analyses of all samples from different size classes showed that oocytes were mature in stages III to IV (Szmant-Froelich et al. 1985; Shikina et al. 2013). Hence, regardless of colony sizes, the reproductive timing of *F. abdita* is similar. This result supports the observations of Maboloc et al. (2016) for April-May samples of the same species from the same region of the Philippines. However, given that the sample size is limited, future studies should involve additional surveys and more samples of colonies across different size classes.

Findings of this study provided information on the size and age at the onset of reproductive maturity and fecundity of sexually produced and wild *F. abdita*. Our results will also be helpful in informing restoration efforts, as determining when sexually cultured and transplanted colonies start to reproduce is important in goal setting during restoration planning. It is recommended, however, to investigate the size at the first maturity of corals with different lifeforms and reproductive strategies to provide a holistic understanding of the population dynamics in the reef.

## Authors’ contributions

**Katya G. Bonilla:** Conceptualization, Methodology, Validation, Formal Analysis, Investigation, Writing; **James R. Guest:** Conceptualization, Methodology, Validation, Investigation, Writing; **Dexter dela W. Cruz:** Conceptualization, Methodology, Validation, Investigation, Writing; **Maria Vanessa Baria-Rodriguez:** Conceptualization, Methodology, Validation, Investigation, Writing.

## Acknowledgments

We would like to thank Toh Tai Chong, Marcos Ponce, Ronald de Guzman, Jun Castrence, Christine Baran, and Jerry Arboleda for their valuable assistance during field and laboratory works and Darryl Valino for the site map.

## References

Adjeroud, M., Kayal, M., Iborra-Cantonnet, C., Bosserelle, P., Liao, V., Chancerelle, Y., Caludet, J., Penim C., 2018. Recovery of coral assemblages despite acute and recurrent disturbances on a South-Central Pacific reef. Sci. Reps. 1–3.

Babcock, R.C., 1984. Reproduction and distribution of two species of *Goniastrea* (Scleractinia) from the Great Barrier Reef Province. Coral Reefs. 187–195.

Babcock, R.C., 1986. Population Biology of Reef Flat Corals of the Family Faviidae *(Goniastrea, Platygyra)*. Thesis. James Cook University of North Queensland, Townsville. 163 pp.

Baird, A.H., Guest, J.R., Willis, B.L., 2009. Systematic and biogeographical patterns in the reproductive biology of scleractinian corals. Annu. Rev. Ecol. Evol. Syst. 551–571.

Bak, R.P.M., Engel, M.S., 1979. Distribution, abundance, and survival of juvenile hermatypic corals (Scleractinia) and the importance of life history strategies in the parent coral community. Mar. Biol. 341–352.

Bak, R.P.M, Meesters, E.H., 1999. Population structure as a response of coral communities to global change. Am. Zool. 56–65.

Baria, M.V.B., dela Cruz, D.W., Villanueva, R.D., Guest, J.R., 2012. Spawning of three-year-old *Acropora millepora* corals reared from larvae in Northwestern Philippines. Bull. Mar. Sci. 61–62.

Bell, G., 1980. The costs of reproduction and their consequences. Am. Nat. 45–76.

Connell, J.H., 1973. Population ecology of reef-building corals, in: Jones, O.A., Endean, R. (Eds.), Biology and Geology of Coral Reefs. Academic Press, New York. 205–245.

Darling, E.S., Alvarez-Filip, L., Oliver, T.A., McClanahan, T.R., Côté, I.M., 2012. Evaluating life-history strategies of reef corals from species traits. Ecol. Lett. 1378–1386.

dela Cruz, D.W., Harrison, P. L., 2017. Enhanced larval supply and recruitment can replenish reef corals on degraded reefs. Sci. Rep. 13985.

Eckelbarger, K. J., 1986. Vitellogenic mechanisms and the allocation of energy to offspring in polychaetes. Bull. Mar. Sci. 426–443.

Edmunds, P.J., Carpenter, R.C., 2001. Recovery of *Diadema antillarum* reduces macroalgal cover and increases abundance of juvenile corals on a Caribbean reef. PNAS. 5067–5071.

Edmunds, P.J., 2007. Evidence for a decadal-scale decline in the growth rates of juvenile scleractinian corals. Mar. Eol. Prog. Ser. 1–13.

Glynn, P. W., Gassman, M.J., Eakin, C.M., Cortes, J., Smith, D.B., Guzman, H.M., 1991. Reef coral reproduction in the eastern Pacific: Costa Rica, Panama, and Galapagos Islands (Ecuador). 1. Pocilloporidae. Mar. Biol. 355–368.

Guest, J.R., Heyward, A.J., Omori, M., Iwao, K., Morse, A., Boch, C., Edwards, A.J., 2010. Rearing coral larvae for reef rehabilitation. In: Edwards, A.J. (Ed.), Reef Rehabilitation Manual. Coral Reef Targeted Research & Capacity Building for Management Program, St. Lucia. 73–92.

Guest, J.R., Baria, M.V., Gomez, E.D., Heyward, A.J., Edwards, A.J., 2014. Closing the circle: is it feasible to rehabilitate reefs with sexually propagated corals? Coral Reefs. 45–55.

Hall, V.R., Hughes, T.P., 1996. Reproductive strategies of modular organisms: Comparative studies of reef-building corals. Ecology. 950–963.

Harrison, P.L., Wallace, C.C., 1990. Reproduction, dispersal, and recruitment of scleractinian corals, in: Dubinsky, Z. (Ed.), Ecosystems of the world, Coral Reefs. 133–207.

Harrison, P., 2011. Sexual Reproduction of Scleractinian Corals. Coral Reefs. 59–84.

Iwao, K., Omori, M., Taniguchi, H., Tamura, M., 2010. Transplanted *Acropora tenuis* (Dana) spawned first in their life 4 years after culture from eggs. Galaxea. 47.

Kai, S., Sakai, K., 2008. Effect of colony size and age on resource allocation between growth and reproduction in the corals *Goniastrea aspera* and *Favites chinensis*. Mar. Ecol. Prog. Ser. 133–139.

Kojis, B.L., Quinn, N.J., 1982. Reproductive ecology of two Faviid corals (Coelenterata: Scleractinia). Mar Ecol. Prog. Ser. 251–255.

Kojis, B.L., and Quinn, N.J., 1985. Puberty in *Goniastrea favulus:* age or size limited? Proceedings of the Fifth International Coral Reef Congress, Tahiti. 289–294.

Kojis, B.L., 1986. Sexual Reproduction in *Acropora (Isopora)* species (Coelenterata: Scleractinia) I. *A. cuneata* and *A. palifera* on Heron Island reef, Great Barrier Reef. Mar. Biol. 291–309.

Llodra, E.R., 2002. Fecundity and life-history strategies in marine invertebrates. Adv. Mar. Biol. 87–170.

Lukoschek, V., Cross, P., Torda, G., Zimmerman, R., Willis, B.L., 2013. The importance of coral larval recruitment for the recovery of reefs impacted by cyclone Yasi in the central Great Barrier Reef. PLoS ONE. e65363.

Maboloc, E.A., Jamodiong, E.A., Villanueva, R.D., 2016. Reproductive biology and larval development of the scleractinian corals *Favites colemani* and *F. abdita* (Faviidae) in northwestern Philippines. Invertebr. Reprod. Dev. 1–11.

McGinley, M. A., Temme, D. H. and Geber, M. A. 1987. Parental investment in offspring in variable environments: theoretical and empirical considerations. American Naturalist 370–398.

Miller KJ, Mundy, C.N., 2003. Rapid settlement in broadcast spawning corals: implications for larval dispersal. Coral Reefs. 99–106.

Moberg, F., Folke, C., 1999. Ecological goods and services of coral reef ecosystems. Ecol. Econ. 215–233.

Nishikawa, A., Katoh, M., Sakai, K., 2003. Larval settlement rates and gene flow of broadcast-spawning *(Acropora tenuis)* and planula brooding *(Stylophora pistillata)*corals. Mar. Ecol. Prog. Ser. 87–97.

Sakai, K., 1998a. Delayed maturation in the colonies coral *Goniastrea aspera* (Scleractinia): whole-colony mortality, colony growth and polyp egg production. Res. Popul. Ecol. 287–292.

Sakai, K., 1998b. Effect of colony size, polyp size, and budding mode on egg production in a colonial coral. Biol. Bull. 319–325.

Schmidt-Roach, S., Miller, K., Woolsey, E., Gerlach, G., Baird, A., 2012. Broadcast spawning by *Pocillopora* species on the Great Barrier Reef. PLoS ONE. e50847.

Shikina, S., Chen, C.J., Chung, Y.J., Shao, Z.F., Liou, J.Y., Tseng, H.P., Lee, Y.H., Chang, C.F., 2013. Yolk formation in a stony coral *Euphyllia ancora* (Cnidaria, Anthozoa): insight into the evolution of vitellogenesis in nonbilaterian animals. Endocrinology. 3447–3459.

Soong, K., 1991. Sexual reproductive patterns of shallow-water reef corals in Panama. Bull. Mar. Sci. 832–846.

Soong, K., 1993. Colony size as a species character in massive reef corals. Coral Reefs. 77–83.

Szmant-Froelich, A., Reutter, M., Riggs, L., 1985. Sexual reproduction of *Favia fragum* (Esper): lunar patterns of gametogenesis, embryogenesis, embryogenesis and planulation in Puerto Rico. Bull. Mar. Sci. 880–892.

Szmant, A.M., 1986. Reproductive ecology of Caribbean reef corals. Coral Reefs. 543–553.

Toh, T.C., Guest, J., Chou, L.M., 2012. Coral larval rearing in Singapore: Observations on spawning timing, larval development, and settlement of two common scleractinian coral species, in: Tan, K.S. (Ed.), Contributions to Marine Science. National University of Singapore, Republic of Singapore, 81–87.

Toh, T.C., 2014. The use of sexually propagated scleractinian corals for reef restoration (Ph.D. Dissertation). National University of Singapore, Singapore.

van Moorsel, G.W.N.M., 1988. Early maximum growth of stony corals (Scleractinian) after settlement of artificial substrata on a Caribbean reef. Mar. Ecol. Prog. Ser. 127–135.

Villanueva, R.D., Yap, H.T., Montano, M.N.E., 2008. Timing of planulation by pocilloporid corals in the northwestern Philippines. Mar. Ecol. Prog. Ser. 111–119.

Wallace, C.C., 1985. Reproduction, recruitment, and fragmentation in nine sympatric species of the coral genus *Acropora*. Mar. Biol. 217–233.

